# Analysis of publicly available transcriptomic data to identify key genes and pathways associated with osteosarcoma metastasis

**DOI:** 10.1101/2024.11.27.623785

**Authors:** Iryna Horak

## Abstract

Osteosarcoma is the most common bone tumor occurring in children and adolescents. The prognosis of osteosarcoma patients with metastasis is rather poor, availability of prognostic molecular markers would thereby help to distinguish patients with a worse prognosis and to choose appropriate treatment. This study aimed to analyze data from publicly available datasets to identify genes and pathways associated with osteosarcoma onset and metastasis.

A total of 8 datasets were analyzed (TARGET-OS, GSE220538, GSE21257, GSE9508, GSE87624, GSE14359, GSE19276, and GSE36001), and common deregulated genes and abundant pathways were searched. Three downregulated genes, *TMBIM4, PKIB* and *IGKC*, were common between metastatic and non-metastatic osteosarcoma tumors. Several abundant GO terms and pathways were identified, including Apoptotic Process (GO:0006915), Regulation Of Phosphatidylinositol 3-Kinase Signaling (GO:0014066), Regulation Of Cell Adhesion Molecule Production (GO:0060353), Positive Regulation Of MAP Kinase Activity (GO:0043406), and KEGG pathway Adherens junction.

Analysis of metastasis *versus* primary tumor revealed 231 common deregulated genes, identified hub genes involved in the organization of cell-cell junctions and surfactant metabolism. Significant enrichment was found in tight junctions, actin cytoskeleton, focal adhesion, muscle contraction proteins, NF-κB, PIK3/Akt/mTOR, AMPK, TNF, and MAPK signaling.

335 common deregulated genes were found between tumor and normal bone, network analysis revealed two clusters involved in cell cycle progression and G2/M transition, and immune response regulation. Abundance was found in p53, TNF, MAPK, and JAK-STAT pathways.

Taken together, this study consolidated transcriptomic data from 8 publicly available datasets to identify common deregulated genes and pathways in osteosarcoma development and metastasis.

## Introduction

Osteosarcoma (OSA) is the most common bone tumor affecting predominantly children and adolescents with approximately 4.4 cases per million children annually [1]. 15-20% of OSA patients were diagnosed with metastasis, and in 85% of all cases OSA metastasizes to lungs, rarely to other bones. Treatment strategy includes surgical removal of the primary tumor and chemotherapy with methotrexate, doxorubicin, and cisplatin [2,3]. Improvement in diagnostics and treatment of OSA led to an increase in 5-year overall survival from less than 20% to 65-70% [2]. However, patients with metastasis still have poor long-term prognosis, around 20-30% [4]. Therefore, molecular markers of OSA motility and metastasis are intensively investigated.

Transcriptomic data may elucidate important genes and pathways involved in OSA onset and metastasis. In this study, 8 datasets containing transcriptomic data from OSA patients were analyzed, and transcriptomic data from OSA tumors *versus* normal bone, primary tumors *versus* metastasis, and metastatic *versus* non-metastatic OSA were compared. GO, KEGG, and WikiPathways enrichment and protein-protein interactions analysis were performed. Key hub genes and signaling pathways associated with OSA development and metastasis were identified.

## Materials and methods

### Data acquisition

Raw RNA-seq counts for TARGET-OS (https://www.cancer.gov/ccg/research/genome-sequencing/target/studied-cancers/osteosarcoma) dataset were obtained from UCSC Xena Browser https://xenabrowser.net/datapages/?dataset=TARGET-OS.star_counts.tsv&host=https%3A%2F%2Fgdc.xenahubs.net&removeHub=https%3A%2F%2Fxena.treehouse.gi.ucsc.edu%3A443. 87 samples with information about metastasis status were available, among them 32 patients diagnosed with metastasis, and 55 – without. There were 50 males and 37 females, and the median age was 14 years.

Other datasets (GSE220538, GSE21257, GSE9508, GSE87624, GSE14359, GSE19276, and GSE36001) were obtained from the NCBI GEO platform, differential expression analysis performed with the GEO2R tool.

### Differential expression analysis of TARGET-OS dataset

For differential expression analysis of TARGET-OS dataset, DESeq2 [5] was utilized. Metastasis group was compared to No Metastasis group, significance cut-offs were padj < 0.05 and |log2FoldChange| > 0.59. Ggplot2, ggplotly, pheatmap, and EnhancedVolcano were used for results visualization.

### Functional analysis

Gene enrichment analysis was performed with enrichR and ClusterProfiler [6,7]. Databases GO (Gene ontology), KEGG (Kyoto Encyclopedia of Genes and Genomes), and WikiPathways were used for enrichment analysis. For GSEA, rank was calculated as sign(log2FoldChange) * (-log10(pvalue)), top-5000 genes arranged by adjusted P value were used for analysis. Overlaps between genes and enriched terms were searched using Venn diagram online tool (https://bioinformatics.psb.ugent.be/webtools/Venn/). Complete results of enrichment analysis are shown in Supplements. For visualization of protein-protein interactions STRING database (https://string-db.org/) was used (minimum interaction score 0.700), and hub genes were identified in Cytoscape software with CytoHubba plugin according to the Degree value.

## Results

### Differential expression and functional analysis of TARGET-OS dataset

Two groups of primary tumors were compared, from patients who developed metastasis (Metastasis), and patients who did not develop metastasis (No Metastasis). Differential expression analysis was performed with DESeq2, and 223 differentially expressed genes were found, among them 99 upregulated in Metastasis group, and 124 downregulated (Fig. 1A-B, interactive volcano plot and differential expression analysis table are available in Supplements). Protein-protein interactions among DEGs were visualized and top-10 hub genes were identified, namely *MYL2, TP53, NEB, KLHL41, ACTA1, MYOZ1, TCAP, KLHL40, NRAP, TNNC2*. Two clusters were found, one involved in myofibril assembly and muscle cell development, and second one, p53-centered, involved in the regulation of apoptosis and cell proliferation (Fig. 1B). DEGs enrichment against GO, KEGG, and WP (WikiPathways) was performed with enrichR package. Selected terms are shown at Fig. 1C. Substantial enrichment in genes associated with muscle contraction and actomyosin complex was observed. In addition, terms related to cell cycle, apoptosis, adhesion, and signaling pathways BMP, Wnt, and p53 were enriched. GSEA analysis was performed with clusterProfiler, and selected deregulated terms are presented at Fig. 1D. Actomyosin structure and cytoskeleton organization, mitochondrial translation, RNA processing, DNA replication, ATP synthesis, and TGFβ signaling were found to be upregulated in primary tumors of patients with metastasis. Cell adhesion, regulation of cell differentiation, cytokine production and immune response, and PI3K/Akt/mTOR signaling were found to be downregulated.

**Fig. 1.**
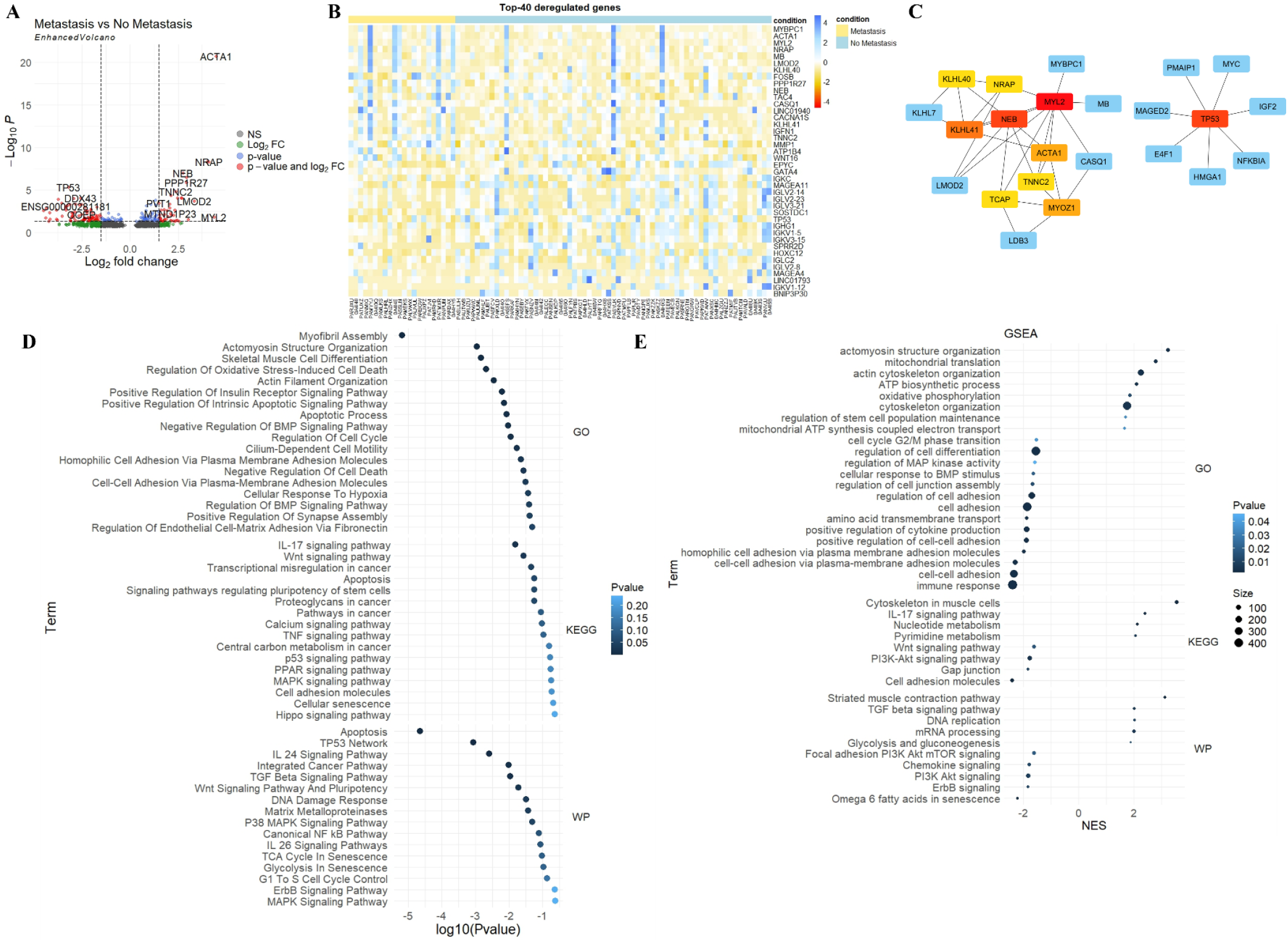
Transcriptomic changes in metastatic OSA: analysis of TARGET-OS dataset. A – volcano plot; B – heatmap representing top-20 upregulated and top-20 downregulated genes; C – network of hub genes; D, E – enrichment analysis and GSEA of deregulated genes (selected terms).

### Analysis of primary tumors

To compare results from TARGET-OS dataset to other OSA data, three publicly available datasets were used: GSE220538, GSE21257, and GSE9508. DE data for these datasets were obtained from GEO2R tool, in each dataset primary tumors from patients with diagnosed metastasis were compared to those without metastasis. Enrichment of obtained DEGs was performed as described before, and overlaps in enriched terms in at least two datasets were sought. Dataset GSE9508 demonstrated a low number of DEGs, so it was not used for enrichment analysis.

Genes *TMBIM4* and *PKIB* were downregulated in datasets TARGET-OS and GSE21257, and *IGKC* downregulation was common for TARGET-OS and GSE9508 datasets. According to enrichment results, 21 GO terms were common for two datasets, including Apoptotic Process (GO:0006915), Negative Regulation Of Glucose Import (GO:0046325), Regulation Of RNA Biosynthetic Process (GO:2001141), Regulation Of Phosphatidylinositol 3-Kinase Signaling (GO:0014066), Regulation Of Cell Adhesion Molecule Production (GO:0060353), Regulation Of Monooxygenase Activity (GO:0032768), and Positive Regulation Of MAP Kinase Activity (GO:0043406). 4 KEGG terms overlapped between two datasets, including terms Bladder cancer and Adherens junction. 6 WP terms were common for two datasets, including IL 24 Signaling Pathway WP5413, Bladder Cancer WP2828, and LTF Danger Signal Response Pathway WP4478.

### Analysis of metastasis vs primary tumor

To compare changes in metastasis versus primary OSA tumors, datasets GSE87624, GSE220538, and GSE14359 were used. 38 common for three datasets upregulated genes were found, and 156 more genes overlapped between the two datasets. *MAGEB2* was downregulated in all datasets, and 36 more genes were downregulated in two datasets. Genes common for at least two datasets were visualized as a network using STRING, they appear to create two main clusters, one involved in tight junction assembly, and second – in surfactant homeostasis. Top-10 hub genes were identified, namely *SFTPA1, SFTPB, SFTPC, OCLN, NKX2-1, CLDN7, TJP3, CDH1, MUC1*, and *SFTPD*, which participate in cell-cell junctions organization and surfactant metabolism (Fig. 2A).

**Fig. 2.**
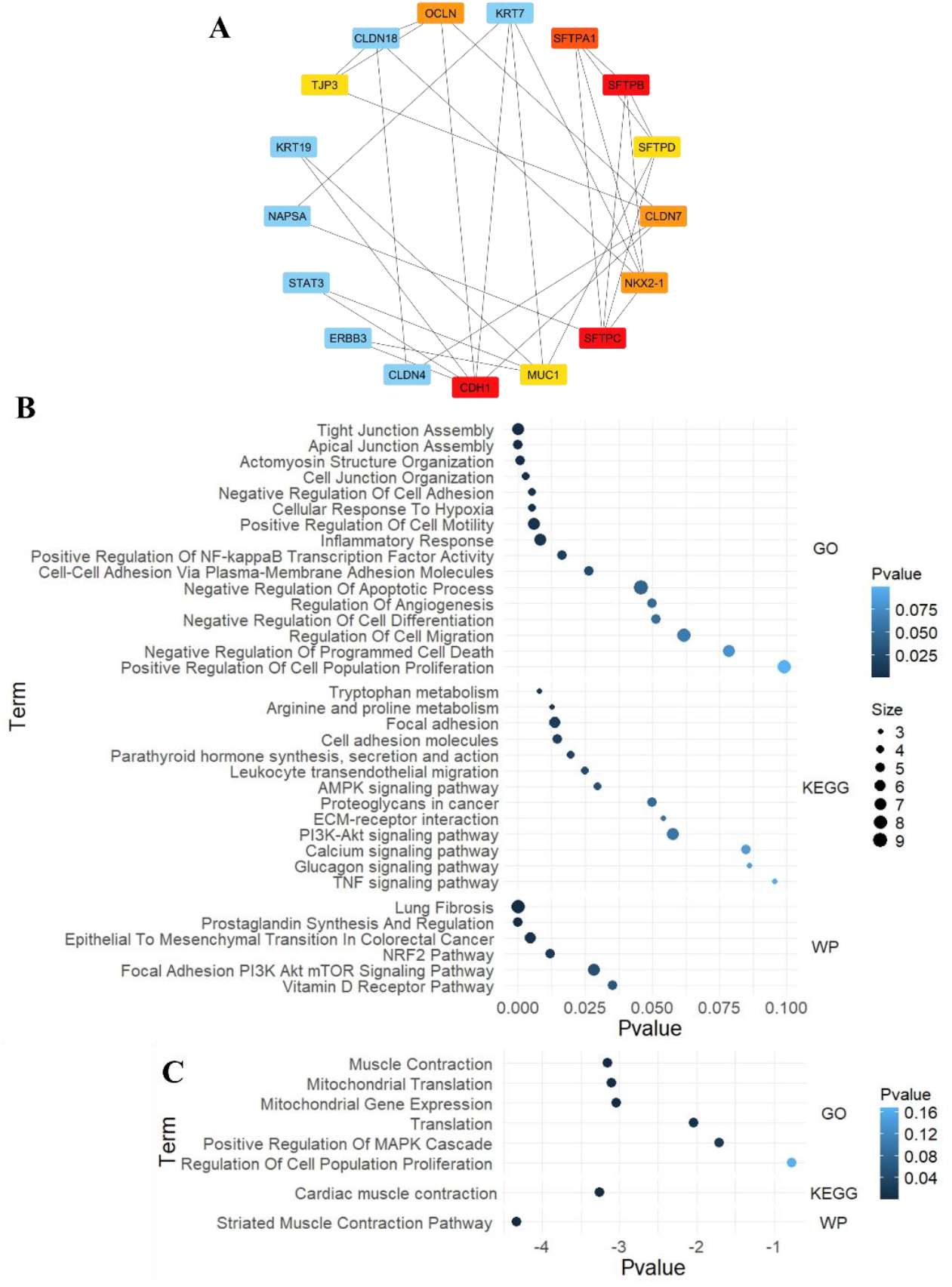
Analysis of deregulated genes in OSA metastasis compared to primary tumor. A – network of hub genes; B, C – enrichment analysis of upregulated and downregulated in metastasis genes (selected terms).

Enrichment analysis of common upregulated genes revealed terms related to tight junctions, actin cytoskeleton, focal adhesion, cell motility, cell death, inflammation, amino acids metabolism, and response to hypoxia. Several signaling pathways were enriched, including NF-κB, PIK3/Akt/mTOR, AMPK, and TNF (Fig. 2B). Downregulated in metastasis genes are abundant in muscle contraction proteins, mitochondrial gene expression, translation, regulation of proliferation, and MAPK cascade (Fig. 2C).

### Analysis of primary tumors vs normal bone

Three datasets containing expression data of primary OSA tumors and normal bones were analyzed: GSE19276, GSE36001, and GSE87624. *CBS* was upregulated in all datasets; another 90 genes were upregulated in two datasets. Among downregulated genes 11 were common for all three datasets (*SORL1, FCN1, ATP8B4, CAT, FGL2, CXCR2, PRKCD, LEPR, FCER1A, DHRS9*, and *TMEM71*), and 233 – for two datasets. A network of common genes revealed two main clusters, one involved in cell cycle progression and G2/M transition, and second – in the regulation of immune response. Top-10 hub genes were identified, all of them engaged in leucocyte activation and regulation of immune response: *LCP2, CD8A, SYK, FCER1G, PECAM1, CD86, FCGR3B, CD2, PTPRC*, and *TYROBP* (Fig. 3A). Upregulated DEGs were enriched in terms related to cell cycle, mitosis, DNA replication and reparation, negative regulation of apoptosis, senescence, amino acids metabolism, and p53 signaling (Fig. 3B). Downregulated genes were abundant in TNF signaling, cytokines production, cell adhesion, regulation of cell migration and apoptosis, MAPK, JAK-STAT, EGFR, VEGF, vitamin D, and NF-κB signaling pathways (Fig. 3C).

**Fig. 3.**
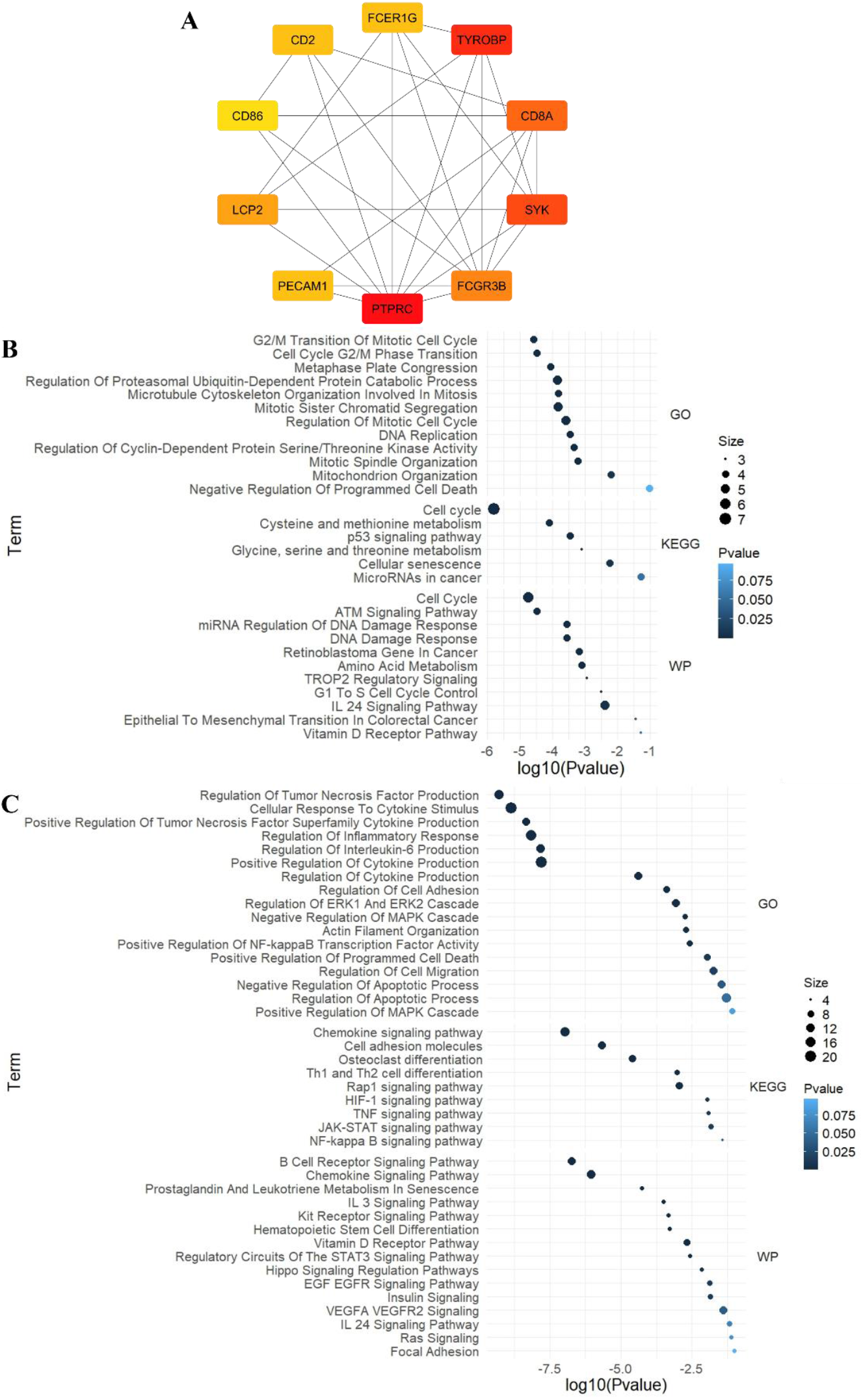
Analysis of deregulated genes in OSA compared to normal bone. A – network of hub genes; B, C – enrichment analysis of upregulated and downregulated in OSA genes (selected terms).

## Discussion

Analysis of TARGET-OS dataset demonstrated that upregulation of cytoskeletal proteins, ATP synthesis, DNA replication, and translation together with downregulated cell adhesion, PI3K/Akt/mTOR signaling involved in focal adhesion, and immune response are associated with OSA metastasis. Hub genes belong to two clusters involved in myofibril assembly and p53-dependent apoptosis and cell proliferation.

Since all four datasets containing data from primary tumors had a relatively small number of DEGs, common enriched terms were analyzed instead. This analysis confirmed the importance of cell adhesion and cell junctions, control of apoptosis as well as PI3K signaling in metastasis.

Between metastatic tissue and primary OSA tumor more DEGs were found, so only common for at least two datasets genes were selected for enrichment analysis. It turned out that DEGs form two clusters involved in tight junction assembly and surfactant homeostasis. Upregulated genes are enriched in cell junctions, cell adhesion, cell motility, negative regulation of apoptosis, NF-κB, NRF, AMPK, and PI3K/Akt signaling, while downregulated genes are enriched in muscle contraction protein, mitochondrial translation, and MAPK signaling.

To identify drivers of osteosarcoma development, DEGs between primary OSA tumors and normal bone were analyzed. It turned out that OSA-associated DEGs are involved in immune response, cell cycle control, p53 signaling, regulation of cell adhesion, migration, and apoptosis.

Experimental data indicate that loss of cell-cell contacts and actin cytoskeleton reorganization are required for OSA cells migration and metastasis [4,8]. Expression of focal adhesion kinase and other focal adhesion proteins as well as PI3K/Akt signaling are associated with OSA migration and metastasis [9–11]. There is evidence that multiple adhesion and cytoskeletal proteins are involved in EMT and facilitate motility and metastasis of OSA cells [12–14]. TGFβ [15–17], NF-κB [12,18,19], MAPK [20,21], HIF-1[22–24], and other signaling pathways [25,26] were demonstrated to participate in OSA progression and metastasis. In addition, enrichment analysis revealed such terms as p53 signaling and control of apoptosis involved in OSA development and metastasis. P53 is known to be mutated in OSA and targeting p53 signaling suppresses tumor growth and mediates sensitivity to chemotherapy by inducing cell cycle arrest, mitochondrial dysfunction, and apoptosis [27–30].

Among hub genes deregulated in OSA metastasis vs primary tumor, CDH1 (E-cadherin) was identified as the one with the highest Degree score. *CDH1* was upregulated in all three analyzed datasets. It is known as a molecular marker of EMT and its downregulation is associated with elevated motility and metastasis of cancer cells [31]. However, there is an opposite process known as mesenchymal-to-epithelial transition (MET) occurring in metastasis, associated with re-expression of *CDH1* and retrieval of adhesion and proliferation [32,33]. Other hub genes, particularly *SFTPB* and *MUC1* are associated with metastasis and poor prognosis in OSA [34,35].

Deregulated between normal bone and OSA genes were abundant in immune response, cell cycle, p53, TNF, and MAPK signaling. Some hub genes identified in this study, namely *LCP2, CD8A, CD2*, and *SYK*, were also identified as potential OSA biomarkers in other bioinformatics papers [36–39]. Moreover, tyrosine kinase SYK is required for adhesion and migration of OSA [40,41]. There is multiple evidence of the involvement of TYROBP in OSA progression, and its association with immune infiltration and better outcome [42–44]. CD86 +1057G/A polymorphism was demonstrated to be associated with elevated susceptibility to OSA [45]. Also, CD86 participates in the interaction between OSA and dendritic cells [46]. Finally, PTPRC, which has the highest degree level in the PPI network, was demonstrated to block OSA cells ferroptosis via regulating FTH1/FTL signaling and to promote autophagy [47].

To summarize, in this study transcriptomic data from 8 publicly available datasets were analyzed to identify common deregulated genes and signaling pathways involved in osteosarcoma development and metastasis. Future validation of obtained results is needed.

## Acknowledgements

Sincere gratitude is expressed to Genomics UA team for conducting the course “RNA-seq: data analysis in R” and providing education in bioinformatics.

This project has received funding through the MSCA4Ukraine project, which is funded by the European Union. Views and opinions expressed are however those of the author(s) only and do not necessarily reflect those of the European Union. Neither the European Union nor the MSCA4Ukraine Consortium as a whole nor any individual member institutions of the MSCA4Ukraine Consortium can be held responsible for them.

## References

[1] Eaton BR, Schwarz R, Vatner R, Yeh B, Claude L, Indelicato DJ, et al. Osteosarcoma. Pediatr Blood Cancer 2021;68. 10.1002/pbc.28352.

[2] Isakoff MS, Bielack SS, Meltzer P, Gorlick R. Osteosarcoma: Current Treatment and a Collaborative Pathway to Success. J Clin Oncol 2015;33:3029–35. 10.1200/JCO.2014.59.4895.

[3] Belayneh R, Fourman MS, Bhogal S, Weiss KR. Update on Osteosarcoma. Curr Oncol Rep 2021;23:71. 10.1007/s11912-021-01053-7.

[4] Sosa-García B, Gunduz V, Vázquez-Rivera V, Cress WD, Wright G, Bian H, et al. A Role for the Retinoblastoma Protein As a Regulator of Mouse Osteoblast Cell Adhesion: Implications for Osteogenesis and Osteosarcoma Formation. PLoS One 2010;5:e13954. 10.1371/journal.pone.0013954.

[5] Love MI, Huber W, Anders S. Moderated estimation of fold change and dispersion for RNA-seq data with DESeq2. Genome Biol 2014;15:550. 10.1186/s13059-014-0550-8.

[6] Kuleshov M V., Jones MR, Rouillard AD, Fernandez NF, Duan Q, Wang Z, et al. Enrichr: a comprehensive gene set enrichment analysis web server 2016 update. Nucleic Acids Res 2016;44:W90–7. 10.1093/nar/gkw377.

[7] Xu S, Hu E, Cai Y, Xie Z, Luo X, Zhan L, et al. Using clusterProfiler to characterize multiomics data. Nat Protoc 2024. 10.1038/s41596-024-01020-z.

[8] Zucchini C, Manara MC, Pinca RS, De Sanctis P, Guerzoni C, Sciandra M, et al. CD99 suppresses osteosarcoma cell migration through inhibition of ROCK2 activity. Oncogene 2014;33:1912–21. 10.1038/onc.2013.152.

[9] Wei Z, Xia K, Zheng D, Gong C, Guo W. RILP inhibits tumor progression in osteosarcoma via Grb10-mediated inhibition of the PI3K/AKT/mTOR pathway. Mol Med 2023;29:133. 10.1186/s10020-023-00722-6.

[10] Cheng S, Liu S, Chen B, Du C, Xiao P, Luo X, et al. Psoralidin inhibits osteosarcoma growth and metastasis by downregulating ITGB1 expression via the FAK and PI3K/Akt signaling pathways. Chin Med 2023;18:34. 10.1186/s13020-023-00740-w.

[11] Chen J-K, Peng S-F, Lai KC, Liu H-C, Huang Y-P, Lin C-C, et al. Fisetin Suppresses Human Osteosarcoma U-2 OS Cell Migration and Invasion via Affecting FAK, uPA and NF-ĸB Signaling Pathway In Vitro. In Vivo (Brooklyn) 2019;33:801–10. 10.21873/invivo.11542.

[12] Liu P, Yang P, Zhang Z, Liu M, Hu S. Ezrin/NF-κB Pathway Regulates EGF-induced Epithelial-Mesenchymal Transition (EMT), Metastasis, and Progression of Osteosarcoma. Med Sci Monit 2018;24:2098–108. 10.12659/MSM.906945.

[13] Shao S, Piao L, Wang J, Guo L, Wang J, Wang L, et al. Tspan9 Induces EMT and Promotes Osteosarcoma Metastasis via Activating FAK-Ras-ERK1/2 Pathway. Front Oncol 2022;12. 10.3389/fonc.2022.774988.

[14] Shao S, Piao L, Guo L, Wang J, Wang L, Wang J, et al. Tetraspanin 7 promotes osteosarcoma cell invasion and metastasis by inducing EMT and activating the FAK-Src-Ras-ERK1/2 signaling pathway. Cancer Cell Int 2022;22:183. 10.1186/s12935-022-02591-1.

[15] Shi Q, Xu J, Chen C, Hu X, Wang B, Zeng F, et al. Direct contact between tumor cells and platelets initiates a FAK-dependent F3/TGF-β positive feedback loop that promotes tumor progression and EMT in osteosarcoma. Cancer Lett 2024;591:216902. 10.1016/j.canlet.2024.216902.

[16] Han Y-L, Luo D, Habaxi K, Tayierjiang J, Zhao W, Wang W, et al. COL5A2 Inhibits the TGF-β and Wnt/β-Catenin Signaling Pathways to Inhibit the Invasion and Metastasis of Osteosarcoma. Front Oncol 2022;12. 10.3389/fonc.2022.813809.

[17] Morice S, Danieau G, Tesfaye R, Mullard M, Brion R, Dupuy M, et al. Involvement of the TGF-β Signaling Pathway in the Development of YAP-Driven Osteosarcoma Lung Metastasis. Front Oncol 2021;11. 10.3389/fonc.2021.765711.

[18] Takeda T, Tsubaki M, Genno S, Tomita K, Nishida S. RANK/RANKL axis promotes migration, invasion, and metastasis of osteosarcoma via activating NF-κB pathway. Exp Cell Res 2024;436:113978. 10.1016/j.yexcr.2024.113978.

[19] Li R, Shi Y, Zhao S, Shi T, Zhang G. NF-κB signaling and integrin-β1 inhibition attenuates osteosarcoma metastasis via increased cell apoptosis. Int J Biol Macromol 2019;123:1035–43. 10.1016/j.ijbiomac.2018.11.003.

[20] Fan M, Zhang G, Chen W, Qi L, Xie M, Zhang Y, et al. Siglec-15 Promotes Tumor Progression in Osteosarcoma via DUSP1/MAPK Pathway. Front Oncol 2021;11. 10.3389/fonc.2021.710689.

[21] Li L, Liang S, Wasylishen AR, Zhang Y, Yang X, Zhou B, et al. PLA2G16 promotes osteosarcoma metastasis and drug resistance via the MAPK pathway. Oncotarget 2016;7:18021–35. 10.18632/oncotarget.7694.

[22] Zhou J, Lan F, Liu M, Wang F, Ning X, Yang H, et al. Hypoxia inducible factor-1α as a potential therapeutic target for osteosarcoma metastasis. Front Pharmacol 2024;15. 10.3389/fphar.2024.1350187.

[23] He G, Nie J-J, Liu X, Ding Z, Luo P, Liu Y, et al. Zinc oxide nanoparticles inhibit osteosarcoma metastasis by downregulating β-catenin via HIF-1α/BNIP3/LC3B-mediated mitophagy pathway. Bioact Mater 2023;19:690–702. 10.1016/j.bioactmat.2022.05.006.

[24] Xu W-N, Yang R-Z, Zheng H-L, Jiang L-S, Jiang S-D. NDUFA4L2 Regulated by HIF-1α Promotes Metastasis and Epithelial–Mesenchymal Transition of Osteosarcoma Cells Through Inhibiting ROS Production. Front Cell Dev Biol 2020;8. 10.3389/fcell.2020.515051.

[25] Lilienthal I, Herold N. Targeting Molecular Mechanisms Underlying Treatment Efficacy and Resistance in Osteosarcoma: A Review of Current and Future Strategies. Int J Mol Sci 2020;21:6885. 10.3390/ijms21186885.

[26] Ji Z, Shen J, Lan Y, Yi Q, Liu H. Targeting signaling pathways in osteosarcoma: Mechanisms and clinical studies. MedComm 2023;4. 10.1002/mco2.308.

[27] Rawat L, Nayak V. Piperlongumine induces ROS mediated apoptosis by transcriptional regulation of SMAD4/P21/P53 genes and synergizes with doxorubicin in osteosarcoma cells. Chem Biol Interact 2022;354:109832. 10.1016/j.cbi.2022.109832.

[28] Kumar A, Kaur S, Dhiman S, Singh PP, Bhatia G, Thakur S, et al. Targeting Akt/NF-κB/p53 Pathway and Apoptosis Inducing Potential of 1,2-Benzenedicarboxylic Acid, Bis (2-Methyl Propyl) Ester Isolated from Onosma bracteata Wall. against Human Osteosarcoma (MG-63) Cells. Molecules 2022;27:3478. 10.3390/molecules27113478.

[29] Yu S, Guo L, Yan B, Yuan Q, Shan L, Zhou L, et al. Tanshinol suppresses osteosarcoma by specifically inducing apoptosis of U2-OS cells through p53-mediated mechanism. J Ethnopharmacol 2022;292:115214. 10.1016/j.jep.2022.115214.

[30] Han G, Zhang Y, Liu T, Li J, Li H. The anti-osteosarcoma effect from panax notoginseng saponins by inhibiting the G 0 / G 1 phase in the cell cycle and affecting p53-mediated autophagy and mitochondrial apoptosis. J Cancer 2021;12:6383–92. 10.7150/jca.54602.

[31] Issagholian L, Tabaie E, Reddy AJ, Ghauri MS, Patel R. Expression of E-cadherin and N-cadherin in Epithelial-to-Mesenchymal Transition of Osteosarcoma: A Systematic Review. Cureus 2023. 10.7759/cureus.49521.

[32] Demirkan B. The Roles of Epithelial-to-Mesenchymal Transition (EMT) and Mesenchymal-to-Epithelial Transition (MET) in Breast Cancer Bone Metastasis: Potential Targets for Prevention and Treatment. J Clin Med 2013;2:264–82. 10.3390/jcm2040264.

[33] Lu W, Kang Y. Epithelial-Mesenchymal Plasticity in Cancer Progression and Metastasis. Dev Cell 2019;49:361–74. 10.1016/j.devcel.2019.04.010.

[34] Feng S, Fu D, Zhang Y, Zhang L, Ji Y, Li H, et al. Serum pro-surfactant protein B is correlated with clinical properties in osteosarcoma patients. Biochem Cell Biol 2023;101:456–63. 10.1139/bcb-2022-0179.

[35] Liu J, Xu Y, Xu T, Liu Y, Liu J, Chai J, et al. MUC1 promotes cancer stemness and predicts poor prognosis in osteosarcoma. Pathol - Res Pract 2023;242:154329. 10.1016/j.prp.2023.154329.

[36] Liang J, Chen J, Hua S, Qin Z, Lu J, Lan C. Bioinformatics analysis of the key genes in osteosarcoma metastasis and immune invasion. Transl Pediatr 2022;11:1656–70. 10.21037/tp-22-402.

[37] Ding F, Tian J, Wu J, Han D, Zhao D. Identification of key genes as predictive biomarkers for osteosarcoma metastasis using translational bioinformatics. Cancer Cell Int 2021;21:640. 10.1186/s12935-021-02308-w.

[38] Li G, Lei J, Xu D, Yu W, Bai J, Wu G. Integrative analyses of ferroptosis and immune related biomarkers and the osteosarcoma associated mechanisms. Sci Rep 2023;13:5770. 10.1038/s41598-023-33009-1.

[39] Sun L, Li J, Yan B. Gene expression profiling analysis of osteosarcoma cell lines. Mol Med Rep 2015;12:4266–72. 10.3892/mmr.2015.3958.

[40] Wu W, Wang L, Li S. Hox transcript antisense RNA knockdown inhibits osteosarcoma progression by regulating the phosphoinositide 3-kinase/AKT pathway through the microRNA miR-6888-3p/spleen tyrosine kinase axis. Bioengineered 2022;13:9397–410. 10.1080/21655979.2022.2059614.

[41] Wang T, Xu Y, Liu X, Zeng Y, Liu L. miR-96-5p is the tumor suppressor in osteosarcoma via targeting SYK. Biochem Biophys Res Commun 2021;572:49–56. 10.1016/j.bbrc.2021.07.069.

[42] Liang T, Chen J, Xu G, Zhang Z, Xue J, Zeng H, et al. TYROBP, TLR4 and ITGAM regulated macrophages polarization and immune checkpoints expression in osteosarcoma. Sci Rep 2021;11:19315. 10.1038/s41598-021-98637-x.

[43] Li J, Shi H, Yuan Z, Wu Z, Li H, Liu Y, et al. The role of SPI1-TYROBP-FCER1G network in oncogenesis and prognosis of osteosarcoma, and its association with immune infiltration. BMC Cancer 2022;22:108. 10.1186/s12885-022-09216-w.

[44] Xu H-R, Chen J-J, Shen J-M, Ding W-H, Chen J. TYRO protein tyrosine kinase-binding protein predicts favorable overall survival in osteosarcoma and correlates with antitumor immunity. Medicine (Baltimore) 2022;101:e30878. 10.1097/MD.0000000000030878.

[45] Wang W, Song H, Liu J, Song B, Cao X. CD86 + 1057G/A Polymorphism and Susceptibility to Osteosarcoma. DNA Cell Biol 2011;30:925–9. 10.1089/dna.2011.1211.

[46] Muraro M, Mereuta OM, Saglio F, Carraro F, Berger M, Madon E, et al. Interactions between osteosarcoma cell lines and dendritic cells immune function: An in vitro study. Cell Immunol 2008;253:71–80. 10.1016/j.cellimm.2008.05.002.

[47] Shao Y, Zuo X. PTPRC Inhibits Ferroptosis of Osteosarcoma Cells via Blocking TFEB/FTH1 Signaling. Mol Biotechnol 2024;66:2985–94. 10.1007/s12033-023-00914-9.

